# The unique morphological basis and parallel evolutionary history of personate flowers in *Penstemon*

**DOI:** 10.1101/2024.12.02.626427

**Authors:** Trinity H. Depatie, Carolyn A. Wessinger

## Abstract

**Premise:** Adaptive radiation in ecologically and morphologically diverse plant lineages presents an opportunity to investigate the rapid evolution of novel floral traits. While some types of floral traits, such as flower color, are well-characterized, other types of complex morphologies remain understudied. One example is occluded personate flowers, dorso-ventrally compressed flowers with obstructed floral passageways, which have evolved in multiple genera, but have only been characterized from snapdragon.

**Methods:** Our study examined the morphological basis and evolutionary history of personate flowers in a clade of *Penstemon* species that includes three personate-flowered species. We characterized floral morphology and inferred phylogenomic relationships for 13 species in this group in order to examine the evolutionary history of personate flowers. We used phylogenomic tests for introgression to examine whether personate-flowered lineages have a history of introgression.

**Results:** Unlike the personate flowers of snapdragon, personate flowers in *Penstemon* are produced by deep pleats in the ventral petal tissue that curve the ventral petal surface upwards, obstructing the floral tube opening. Our phylogenetic tree suggests that personate flowers evolved in two separate lineages. Phylogenomic analyses indicate incomplete lineage sorting and introgression between certain taxa have contributed to phylogenomic discordance, however we found little evidence of recent introgression between the two personate-flowered lineages.

**Conclusions:** The evolution of personate flowers in *Penstemon* involves a distinct morphological basis than snapdragon. Personate flowers have evolved multiple times in *Penstemon* on a rapid evolutionary timescale. The source of genetic variation for repeated shifts may be de novo mutations or pre-existing variants.

## INTRODUCTION

Adaptive radiation, the rapid evolution of ecologically diverse species within a lineage, is an important driver of biological diversity (Simpson, 1953; Givnish and Sytsma, 2000). Examples of adaptive radiation are ubiquitous across the tree of life, including many examples described in plants (Kapralov et al., 2013; Breitkopf et al., 2015; Nge et al., 2021; Schenk, 2021). Since Darwin described his observations of the finches on the Galapagos islands (1859), biologists have sought to characterize the features that promote adaptive radiation, including ecological opportunity, patterns of geographic dispersal to new habitats, and genetic mechanisms for phenotypic and species diversification (Schluter, 1996; Losos and Mahler, 2010; Yoder et al., 2010). With the increasing availability of quality genomic resources, an area of considerable current interest is the source of genetic variation that enables the emergence of rapid trait divergence and speciation on a short evolutionary timescale (Walter et al., 2018; Schenk, 2021).

It is now clear that a common feature of adaptive radiation is a signature of phylogenomic discordance, where individual genetic loci exhibit conflicting evolutionary histories (Seehausen, 2004; Wu et al., 2018). Such discordance arises from incomplete lineage sorting (ILS) – when shared ancestral polymorphisms persist within rapidly speciating lineages. When such polymorphisms eventually fix, the resulting pattern of genetic variation will not necessarily reflect the history of speciation events. ILS is prevalent when ancestral populations are large and speciation occurs rapidly (Hudson, 1983; Pamilo and Nei, 1988; Suh et al., 2015; Pease et al., 2016; Alexander et al., 2017). Phylogenomic discordance within adaptive radiations can also arise from genetic introgression among hybridizing incipient species (Schluter, 2000). In this scenario, a hybridization event may transfer alleles across species boundaries. Introgression causes the genealogy of a given locus to conflict with the overall species branching pattern, reflecting relatedness between hybridizing lineages. Hybridization is a common feature of adaptive radiations, particularly in groups with incomplete reproductive barriers and geographic proximity (Seehausen, 2004).

Although ILS and introgression both generate discordance among loci, these processes leave distinct phylogenomic signatures, allowing them to be disentangled (Degnan and Rosenberg, 2009; Hibbins and Hahn, 2021). With ILS, the two possible discordant topologies for a given 3-taxon subtree will occur with equal frequencies across the genome, and divergence times between alleles will be deep, pre-dating the most recent speciation events. With introgression, one of the two discordant topologies will be disproportionately observed in the genome, reflecting relatedness of introgressing lineages. In addition, divergence times between affected sequences will be shallow, reflecting the post-speciation introgression event.

Both processes have the potential to shape patterns of phenotypic evolution within adaptive radiations. If ILS or introgression affects an allele that causes trait divergence, the resulting phylogenetic pattern of trait evolution will be incongruent with the species tree and will potentially suggest convergent evolution (multiple evolutionary transitions), a phenomenon termed ‘hemiplasy’ (Avise and Robinson, 2008; Hibbins et al., 2020). Furthermore, introgression in particular has the potential to enable novel sets of alleles to combine within descendent species, potentially fueling phenotypic diversification. For example, in African rift lake cichlids, hybridization events have generated diversity in physical, behavioral, and ecological traits (Meyer et al., 2019; Urban et al., 2021; Meier et al., 2023). Similarly, in Darwin’s finches, hybridization has reshuffled ancestral haplotypes into novel combinations, generating the observed phenotypic diversity across species (Rubin et al., 2022).

The spectacular diversity in floral form across flowering plants reflects, in part, adaptive radiation within specific lineages (Schenk, 2021). Much attention has been paid to conspicuous floral traits such as flower color, symmetry, floral dimensions, and mating systems. In addition to these well-studied traits, diverse complex floral shapes also have emerged in adaptive radiations. For example, the genus *Aquilegia* (70 species; Munz, 1946) represents an adaptive radiation driven by the evolution of a key floral innovation – nectar spurs (Ree, 2005) – that has likely fueled diversification through pollinator-mediated adaptation and speciation (Hodges and Arnold, 1995). The simple genetic architecture underlying the color and morphology of nectar spurs as well as the colocalization of the genetic loci responsible for these traits (Hodges et al., 2002) likely facilitated *Aquilegia*’s rapid diversification.

Yet many floral morphological traits remain understudied, despite their recurrent evolution. One mysterious floral trait is personate flower shape. Personate flowers are characterized by an upward bulge in the lower petal lobes that fully obstructs or “occludes” the floral passageway (Weberling, 1992). Such flowers are best characterized in snapdragons (*Antirrhinum*), where the personate floral morphology is achieved by a floral hinge formed in the tissue where the upper and lower petal lobes meet (Appendix S3). This morphological innovation apparently acts to protect nectar by filtering visitation to effective types of pollinators (Weberling, 1992). Personate flowers are evolutionarily constrained in snapdragons – all *Antirrhinum* species have this floral type. However, there is variation in this trait across the tribe Antirrhineae (29 genera within family Scrophulariaceae), where four genera contain species-level variation in floral type, categorized into three types: fully occluded (personate), partially occluded, and not occluded (“open”) flowers (Guzmán et al., 2015). Personate flowers were inferred to be the ancestral state in this tribe with an average of 6.91 and 1.5 transitions to partially occluded and open lineages, respectively, suggesting that the loss of personate flowers is evolutionarily labile at the species level (Guzmán et al., 2015). In addition to Antirrhineae, personate flowers have evolved independently in several plant lineages such as *Chelone* and *Penstemon*. This repeated evolution suggests there is likely an adaptive advantage for personate flowers, making this an attractive trait for understanding floral morphological innovation.

The North American wildflower genus, *Penstemon*, includes about 270 species (Wolfe et al., 2006, 2021). Macroevolutionary analyses suggest *Penstemon* exhibits classic patterns of adaptive radiation, with extremely high diversification rates over the past 2.5 million years (Wolfe et al., 2021). *Penstemon*’s geographic origin is inferred to be the Eastern Cordillera and its radiation across the continent has likely involved repeated dispersal through founder events (Wolfe et al., 2006, 2021). Phylogenomic discordance is pervasive in *Penstemon*, likely due to both ILS and introgression, as hybridization is common in the genus (Wilson and Valenzuela, 2002; Cardona et al., 2020; Stone and Wessinger, 2024).

*Penstemon* displays tremendous ecological diversity, including diversification in floral traits. Most species are pollinated by bees, and the genus is well studied for its convergent evolution of hummingbird pollination from bee pollination (Castellanos et al., 2004; Wilson et al., 2006, 2007; Wessinger et al., 2019). However, there is a large amount of diversity within the bee pollinated species that includes variation in flower size, shape, and color. Personate flowers have repeatedly evolved from non-personate ancestors in two *Penstemon* lineages. Subgenus Dasanthera includes two personate species (*P. lyallii* and *P. personatus*) and section Penstemon subsection Penstemon (from here on: subsect. Penstemon) includes three personate species: *P. hirsutus, P. oklahomensis,* and *P. tenuiflorus*.

The latter three species are diploid members of a largely eastern clade within *Penstemon* that is thought to be the most recent expansion within the genus – biogeographic modeling inferred that the clade expanded from the Appalachian Mountains into the southern interior lowlands and coastal plains during the past 1-0.5 MYA (Wolfe et al., 2021). The evolutionary relationships among subsect. Penstemon are especially understudied, and therefore it is unclear whether the three personate species are monophyletic or have evolved in parallel. Wolfe and colleagues (2006 & 2021) inferred a genus-wide phylogeny for *Penstemon*, however, the sampling was incomplete for certain sections, including subsect. Penstemon. In particular, the tree generated by Wolfe and colleagues (2021) was based on 43 nuclear genes and sampled two of the three personate species (*P. hirsutus* and *P. oklahomensis*), finding that they were not sister taxa. At face value, this result suggests that personate flowers evolved multiple times within this clade. However, this tree had fairly low support.

In this study, we characterized the morphological basis of floral shape variation within subsect. Penstemon, estimated phylogenetic relatedness using whole-genome resequencing data for all diploid species of this subsection, and traced the evolutionary history of personate flowers in the group. Additionally, we report a new chromosome-level and annotated de novo genome assembly for *P. smallii*. We found that personate flowers in eastern *Penstemon* species have a different morphological basis than the well-studied personate snapdragon (*Antirrhinum*). Our phylogenomic results find that personate flowers evolved in two lineages in sect. Penstemon, suggesting parallel transitions. However, we identified significant phylogenomic discordance across most internal branches of the phylogeny. Signals of allele sharing between personate lineages appear consistent with ILS, although we can’t rule out a history of introgression. Overall, our results suggest that rapid speciation events, introgression, and geographic dispersal eastwards have shaped the evolutionary history of subsect. Penstemon species.

## MATERIALS AND METHODS

### Study system and sampling

Subsect. Penstemon includes 17 species that vary in ploidy: most are diploid, but there are four known polyploids (*P. calycosus*, *P. deamii*, *P. digitalis*, and *P. laevigatus*) with open flowers that form a monophyletic group (Wolfe et al., 2021). We excluded these four species as well as *P. kralii*, a likely polyploid related to these four. The remaining diploid species exhibit corolla shape variation that includes open, tubular, and personate flowers (Figure 1; Table 1). The three species with personate flowers display flowers that are completely occluded by upwardly curved ventral petal surfaces. Interestingly, the three personate species share additional floral traits: white or pale flower color, lack of nectar guides, and an incredibly hairy yellow staminode (Figure 1D-F; Table 1; Appendix S2), indicating that a suite of correlated traits accompany personate flowers in the eastern subsect. Penstemon clade. By contrast, the species with open (not occluded) corollas have purple flowers with bold nectar guides and display wide openings with sturdy lower lip petals for bees to land on (Figure 1A; Table 1; Appendix S2). The species with tubular flowers have flowers that are only partially occluded by the ventral petal surface and tend to be variable in flower color, nectar guides, and flower shape (Figure 1B-C; Table 1; Appendix S2).

**Figure 1.**
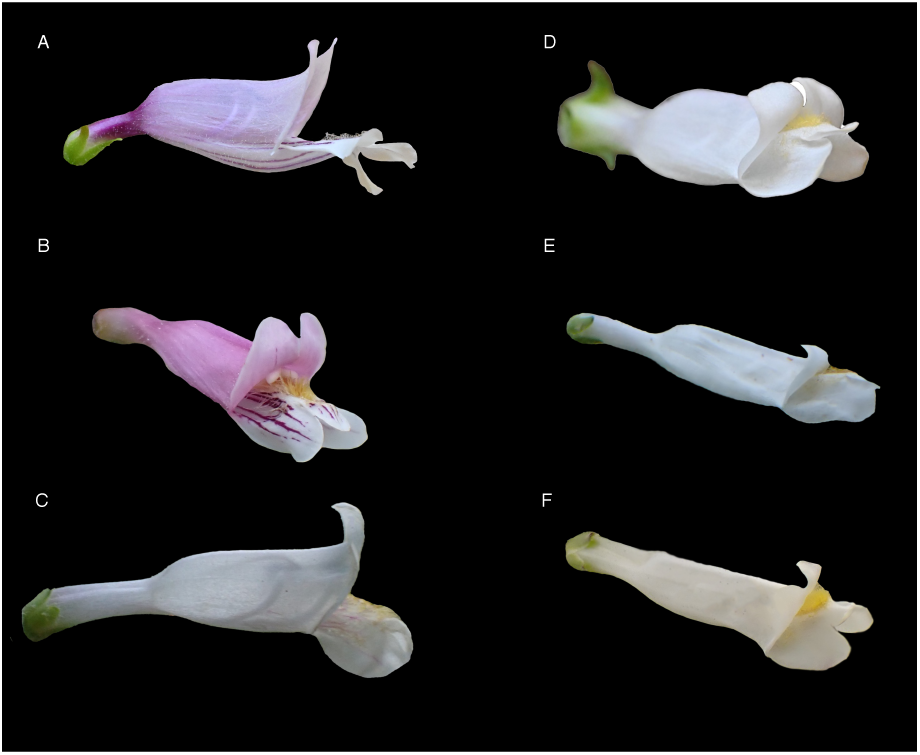
Lateral flower images of open, tubular and personate species in *Penstemon* subsect. Penstemon. (A) *P. smallii,* (B) *P. australis*, (C) *P. pallidus*, (D) *P. hirsutus*, (E) *P. tenuiflorus*, and (F) *P. oklahomensis*.

**Table 1.** An overview of the morphological features of species within *Penstemon* subsect. Penstemon species and clade specificity for species included in the *Twisst* analysis. N/As for specific species indicate omission in morphology analyses or topology weighting. Values of n within morphological analysis columns indicate the number of flowers measured per species.

| Species name | Flower morphology | Average floral obstruction | Average pleat depth | Number of populations re-sequenced | *Twisst* clade designation |
| --- | --- | --- | --- | --- | --- |
| *P. dissectus* | Open | 26.58% (n= 2) | N/A | 1 | N/A |
| *P. smallii* | Open | 26.02% (n= 10) | 15.23% (n=13) | 2 | N/A |
| *P. tenuis* | Open | 24.91% (n= 5) | N/A | 1 | N/A |
| *P. gracilis* | Tubular | N/A | N/A | 1 | N/A |
| *P. canescens* | Tubular | 42.07% (n= 9) | 24.88% (n= 5) | 3 | Eastern Tubular |
| *P. brevisepalus* | Tubular | 37.84% (n= 3) | N/A | 1 | Eastern Tubular |
| *P. australis* | Tubular | 55.17% (n= 14) | 38.58% (n= 5) | 4 | Eastern Tubular |
| *P. laxiflorus* | Tubular | 45.04% (n= 10) | N/A | 2 | N/A |
| *P. arkansanus* | Tubular | 27.72% (n= 7) | N/A | 1 | Western Tubular |
| *P. pallidus* | Tubular | 31.79% (n= 2) | N/A | 2 | Western Tubular |
| *P. hirsutus* | Personate | 78.07% (n= 11) | 46.47% (n= 5) | 3 | Eastern Personate |
| *P. tenuiflorus* | Personate | 76.84% (n= 4) | 45.10% (n= 3) | 2 | Eastern Personate |
| *P. oklahomensis* | Personate | 81.93% (n= 5) | 47.01% (n= 2) | 4 | Western Personate |

We sampled 27 individuals, representing 13 species in subsect. Penstemon (Appendix S1). We included multiple samples for all three species with personate flowers (*P. hirsutus*, *P. oklahomensis*, and *P. tenuiflorus*) as well as five species with open or tubular flowers (*P. australis*, *P. canescens*, *P. laxiflorus*, *P. pallidus,* and *P. smallii*). We included *P. dissectus* (subsection dissecti) as an outgroup.

### Morphological assessment

For all individuals, except *P. arkansanus* and *P. pallidus*, we quantified degree of occlusion from size-standardized photographs of flowers collected in the field. We collected one flower per plant for two to five randomly selected plants within each sampled field population. We measured relative floral occlusion from side-view photographs as the percent of the floral opening height that is blocked by the inward curvature of the ventral petal surface (Figure 2A-C). Because we lacked photos of *P. arkansanus* and *P. pallidus*, we downloaded clear side-view photographs of these species from iNaturalist with observer permission.

**Figure 2.**
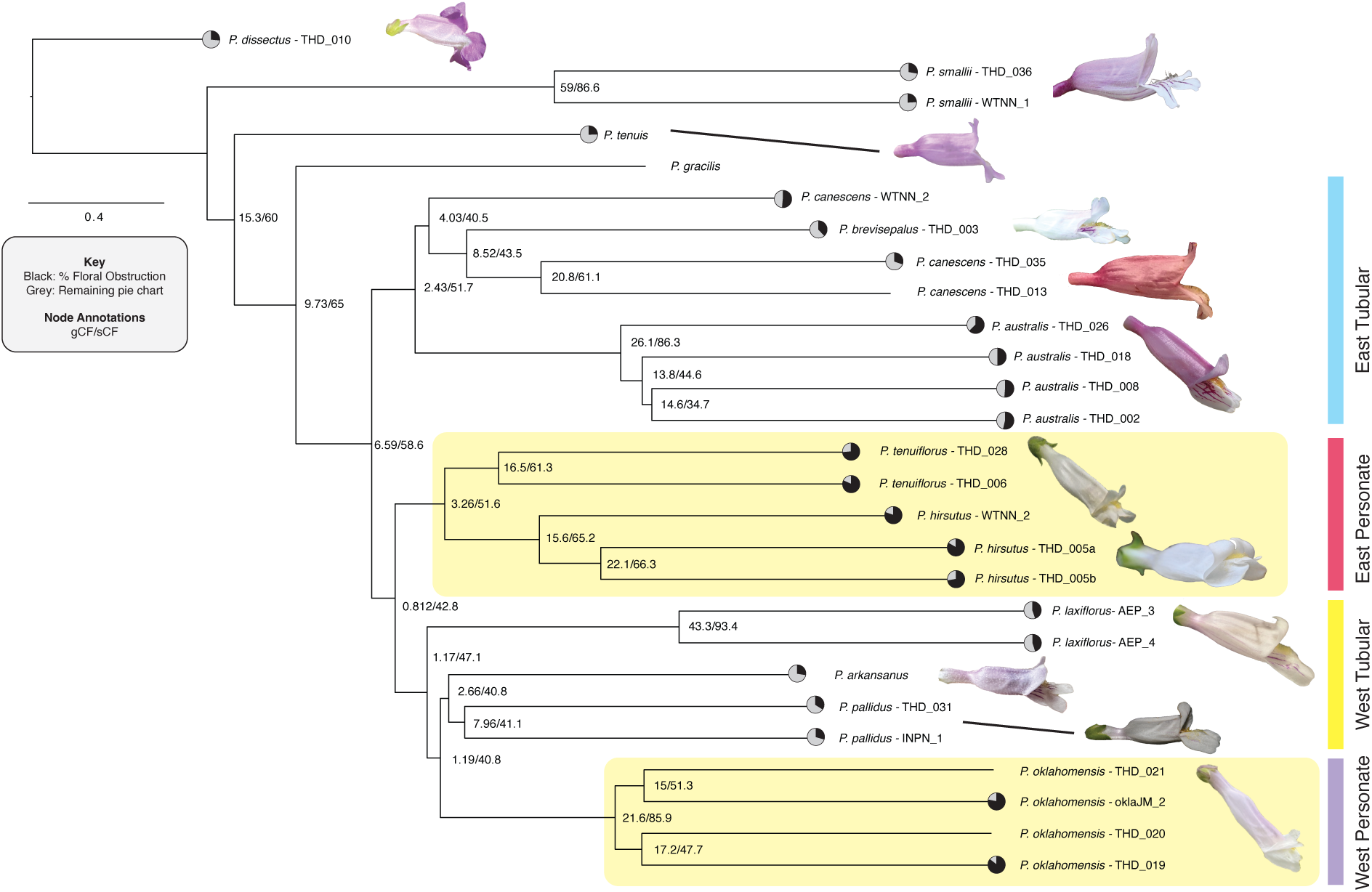
Species tree constructed in Astral-III from CDS sequences for all included subsect. Penstemon species. Black portion of pie charts represent the occlusion of each sample/tip created from floral occlusion measurements. Yellow boxes indicate personate species. Vertical bars specify the species’ clade designation used in analyses. Lateral image of *P. arkansanus* taken by iNaturalist user Luke Benjamin and the photo of *P. pallidus* was taken by iNaturalist user Brian Finzel. All other photos by T. Depatie.

For a subset of populations (see Table 1), we characterized a key feature of *Penstemon*’s personate flowers - the floral tissue on the ventral surface of the corolla tube that folds inward - called “pleats”. These floral pleats produce the occluded floral shape of *Penstemon* flowers by forming the upward bulge in the ventral petal surface. To quantify relative pleat depth, we measured the height of floral pleats relative to the floral tube height in photographs of cross-sections of the floral tube opening just behind the petal lobes (Figure 2A-C). We used the Straight Line tool in ImageJ v2.1.0 (Abràmoff et al., 2004) for all measurements.

### *P. smallii* genome development

A *P. smallii* genome was sequenced, assembled, and annotated by Phase Genomics (Seattle, WA). Leaf tissue was harvested from a *P. smallii* plant obtained from Wood Thrush Native Nursery (Floyd, VA). Genomic DNA was extracted using the TaKaRa NucleoBond HMW DNA kit (TaKaRa Bio, USA). DNA was sheared to a target size of 15-18 kb and purified using AMPure PB beads (Pacific Biosciences). A SMRTbell library was prepared with a mode size of approximately 12 kb and an average size of 18 kb, with minimal fragments below 10 kb. This library was sequenced on a PacBio Sequel II system using two SMRT Cells 8M, generating approximately 990,000 HiFi reads totaling 9 Gb of data. Chromatin conformation capture data was generated using a Phase Genomics Proximo Hi-C 4.0 Kit (Seattle, WA). Briefly, intact cells were crosslinked using a formaldehyde solution, digested using the DPNII, DdeI, HinfI, MseI restriction enzymes, and proximity ligated with biotinylated nucleotides. The resulting Hi-C library was then sequenced on an Illumina NovaSeq generating a total of 113,121,813 150-bp paired-end reads.

Hi-C and PacBio Hifi reads were used in hifiasm (Cheng et al., 2021) with default parameters to generate phased haplotype assembly drafts. Hi-C reads were aligned to the hifiasm draft assemblies, following the Phase Genomics Proximo Hi-C Kit recommendations. Briefly, reads were aligned using bwa-mem (Li, 2013) with the -5SP and -t 8 options specified, and all other options default. SAMBLASTER (Faust and Hall, 2014) was used to flag PCR duplicates for removal. Alignments were then filtered with samtools (Li et al., 2009) using the -F 2304 filtering flag to remove non-primary and secondary alignments. Phase Genomics’ Proximo Hi-C genome scaffolding platform was used to create chromosome-scale scaffolds following the single-phase scaffolding procedure described in Bickhart et al. (2017). Juicebox (Rao et al. 2014; Durand et al. 2016) was used to manually correct scaffolding and assembly errors. The resulting genome assembly included 940 scaffolds covering 438,512,763 bp. The eight largest scaffolds are chromosome-scale, reflecting the haploid chromosome number in *Penstemon*, and cover 404,087,496 bp. We used BUSCO v5.4.4 (Manni et al., 2021) with the eudicots_odb10 database to assess the completeness of our assembly. We found that 97.3% of 2,326 single-copy plant genes are complete, 7.1% are duplicated, and 2.1% are missing.

To annotate the *P. smallii* genome, STAR (Dobin et al., 2013) was used to align RNA-seq data from the closely related *P. barbatus* and *P. kunthii*. Using this as evidence, as well as previous annotations from these species (Wessinger et al., 2023), annotations were generated using GeMoMa v1.9 (Keilwagen et al., 2016, 2018). The annotation includes 25,896 predicted genes.

### DNA extraction, sequencing, and dataset preparation

We extracted DNA from silica-dried leaf tissue using a modified CTAB protocol (Doyle and Doyle, 1987). We performed whole-genome Illumina library preparations and sequencing to ∼10× depth using 150-bp paired-end reads generated on the NovaSeq (Illumina Inc.) platform at Duke University. We used FastP (Chen et al., 2018) to quality filter raw Illumina reads, enabling auto-detection of adapters, limiting read length to 30 bp, filtering out unpaired reads, enabling base correction for overlapping reads, and enabling poly-x trimming on 3′ ends of reads. We then used Fastqc (Andrews, 2010) and multiqc (Ewels et al., 2016) to assess sequence quality. We used bwa-mem to map quality-filtered reads to the *P. smallii* genome and then used bcftools (v1.15.1) (Li et al., 2011) to remove reads with low mapping quality (MQ<30). We marked and removed duplicate reads with samtools markdup and used bamutil clipOverlap (Jun et al., 2015) to clip overlapping paired-end reads. We used bcftools to call genotypes from our filtered reads to produce an “all sites” VCF file that includes both variant and invariant sites.

We used vcftools v0.1.16 (Danecek et al., 2011) to filter variant sites in the all sites VCF to only include sites with maximum individual alleles = 2, minimum site quality of 20, and excluded variants with more than 10% missing data. We estimated genome-wide averages of coverage depth to filter variants for minimum (2), maximum (30) and mean-maximum (20) depths. We then used the filtered all sites VCF file to generate individual gene CDS, genomic window, and consensus whole genome sequences for phylogenetic relationship estimation. We also used vcftools to filter the filtered all sites VCF file for biallelic SNPs. This filtered biallelic SNPs VCF was used as input for tests of introgression (Dsuite and *Twisst* – see below).

### Estimating phylogenetic relationships

We inferred species trees in three ways to confirm patterns of species tree inference: a species tree based on gene coding sequences (CDS) trees, a species tree based on 20 kb genomic window trees, and a concatenated data tree. For the CDS-based species tree, we used gffread v0.12.7 (Pertea and Pertea, 2020) and the annotations from the *P. smallii* reference genome to generate fasta files with spliced exons (CDS) for each of our 27 samples. We then filtered individuals with missing data >50% for each CDS using a custom Python script (Stone and Wessinger, 2024). We estimated gene trees for each CDS in IQ-TREE v2.3.6 (Minh et al., 2020) with the (-mfp) model finder option (Kalyaanamoorthy et al., 2017). We constructed the species tree in ASTRAL-III (Zhang et al., 2018) with the full annotation option (-t 2) to obtain additional quartet information for each branch. For the 20 kb window-based species tree, we split whole-genome sequences into nonoverlapping 20-kb windows and removed windows containing missing data (Ns) more than 75% for any sample using a custom Python script (Stone and Wessinger, 2024). We then used IQ-TREE to estimate “window trees” for each genomic window and used ASTRAL-III to infer the species tree. For the concatenated tree, we used IQ-TREE to estimate a concatenated Maximum Likelihood (ML) tree, by specifying the GTR + I + R substitution model and performing 1,000 ultrafast bootstrap replicates (Hoang et al., 2018). We rooted each tree using the outgroup - *P. dissectus*.

### Quantifying concordance in the CDS species tree

To quantify genealogical concordance across the genome, we used IQ-TREE to compute maximum likelihood gene and site concordance factors (Mo et al., 2023) for each internal branch of the species tree, using CDS sequence alignments and the corresponding CDS-based species tree as our reference tree.

Gene concordance factors (gCFs) reflect the proportion of gene trees that contain the focal branch. Site concordance factors (sCFs) are estimated in quartets and quantify the proportion of sites supporting a particular quartet topology and branch length in the reference species tree. Estimating both gCFs and sCFs can inform on the amount of genealogical concordance of the individual gene trees as well as the informative sites in the sequence data (Lanfear and Hahn, 2024).

### Analysis of trait evolution

We tested whether the degree of floral occlusion differed between floral types (personate, tubular, and open) by performing a phylogenetic ANOVA that corrects for relatedness among species using the phylANOVA function in the R package geiger (Harmon et al., 2008). We also tested whether the degree of floral occlusion is significantly associated with floral pleat depth using a phylogenetic generalized linear model using the glm function in the R package nlme (Pinheiro et al., 2018).

### Identifying genome wide signals of introgression

Using the CDS-based species tree as input, we calculated D and f_4_-ratio statistics for each possible rooted triplet with the program Dsuite v0.5 (Malinsky et al., 2021) and visualized the results with the *f*-branch (*f*_b_) metric. This metric was designed to disentangle correlated f_4_-ratio results and identify excess allele sharing between a particular branch and species combination. Dsuite calculates each statistic with allele frequency estimates, instead of site pattern counts, to allow sampling of multiple individuals for a given population or taxon.

### Assessing local phylogenetic discordance

We used topology weighting (Martin and Van Belleghem, 2017) to quantify patterns of phylogenetic discordance in non-overlapping windows of 100 SNPs across the genome and to identify genomic regions where personate taxa are monophyletic. We first phased our biallelic VCF file to infer haplotypes from SNP genotypes using Beagle v5.4 (Browning et al., 2021). We then used PhyML (Guindon et al., 2010) to infer local neighbor-joining trees from SNPs extracted in non-overlapping 100-SNP windows across the genome. Window trees were inferred under the GTR substitution model. This process yielded 78,057 window trees (mean window size: 2.1kb; Appendix S4).

We next calculated topology weights for the window trees using *Twisst* (Martin and Van Belleghem, 2017). We focused our *Twisst* analysis on the clade containing personate and western tubular species (see Table 1). Specifically, we examined relationships between three clades: eastern personate (*P. hirsutus* and *P. tenuiflorus*), western tubular (*P. arkansanus* and *P. pallidus*), and western personate (*P. oklahomensis*), using the eastern tubular clade (*P. australis*, *P. brevisepalus*, and *P. canescens*) as an outgroup. We calculated topology weights for three possible tree topologies. Topology 1 matches the species tree topology, where the western tubular clade is sister to the western personate clade. Topology 2 is the “miscellaneous” discordant topology where the western tubular clade is sister to the eastern personate clade – this topology is not of specific interest but instead serves as a control topology to help distinguish ILS from introgression. Topology 3 is the “personate” discordant topology where the eastern and western personate clades are monophyletic. Following Stankowski et al. (2024), we utilized a ternary framework to visualize and quantify the distribution of topology weightings across the genome.

### Calculating Relative Node Depth to identify relative divergence between personate taxa

Local trees could show a personate discordant topology due to either ILS or introgression. In theory, these two processes may be distinguished by examining sequence divergence between the two personate lineages, relative to divergence from an outgroup. Divergence times between sequences of the two personate lineages should be relatively deep under ILS (pre-dating the divergence of the western tubular and western personate clades), and shallow under introgression (reflecting recent post-speciation hybridization). We calculated Relative Node Depth (RND) values of 100 SNP windows (used in *Twisst* analysis) to distinguish these possibilities. RND is quantified by calculating genetic distance (*d_XY_*) between two sub-clades relative to their average distance to an outgroup (Hahn, 2018):

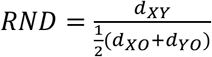

where *X* and *Y* are two focal subclades and *O* is an outgroup. We focused on two RND calculations: *RND_western_* where the focal subclades are the western tubular and western personate clades, and *RND_personate_* where the focal subclades are western personate and eastern personate clades. We used *P. dissectus* as an outgroup for our RND calculations as it likely has minimal history of introgression between the subclades of interest. To calculate these window-based *d_XY_* values, we used our all sites VCF file as input to pixy (Korunes and Samuk, 2021) and calculated *d_XY_* between focal clades in non-overlapping windows of 100 SNPs as used for the *Twisst* analysis.

We performed both RND calculations in two sets of 100 SNP genomic windows, yielding distributions of RND values. First, we calculated both RND values in windows that strongly support the species tree (topology 1 weighting > 0.90), to find the distribution of RND values for windows that follow the species tree as a baseline for comparison. Second, we calculated both RND values for windows that strongly support the personate topology (topology 3 weighting > 0.90), to examine *RND_personate_* values for these discordant windows. We additionally calculated *RND_western_* and *RND_personate_* for a more stringent set of windows: those fully supporting topology 1 compared to those fully supporting topology 3.

### Identifying geographic patterns in population relatedness

To examine geographic patterns of population relatedness in our phylogenomic dataset, we projected a pairwise genetic distance matrix generated using a custom python script (Wessinger et al., 2023) into two dimensions using Multi-Dimensional Scaling (MDS), using the cmdscale function in the R package stats (R Core Team, 2023). We also used Plink v1.90b6.12 (Purcell et al., 2007) to prune SNPs in LD from our filtered all sites VCF and conduct a genomic PCA. Specifically, our LD pruning excluded any SNP that had a r^2^ value greater than 0.10 in windows of 50kb, sliding every 10kb. We examined the relationship between population geographic origin (longitude) and MDS or PCA axis. For this analysis, we focused on personate and tubular species and removed all nursery-derived individuals (n=5) because their geographic origin is unknown.

## RESULTS

### Personate flowers have apparently evolved in parallel within subsect. Penstemon

We created trees using three different approaches – a species tree built with CDS sequences, a species tree built with 20kb genomic windows, and a maximum likelihood concatenated tree. All trees show the same topology, where the three personate species (*P. hirsutus*, *P. tenuiflorus*, and *P. oklahomensis*) are not a monophyletic group (Figure 2; Appendix S5-6). Instead, two personate species (*P. hirsutus* and *P. tenuiflorus*) are sister species, forming an “eastern personate” lineage while the third personate species (*P. oklahomensis*; “western personate”) is more closely related to a “western tubular” clade (*P. pallidus* and *P. arkansanus*). To understand genealogical concordance across our subsect. Penstemon phylogeny, we calculated both gene and site concordance factors on all internal branches of the CDS-based tree. Despite the agreement across our inferred trees, gene and site concordance factors were low, particularly within the clade of personate and tubular species, reflecting substantial genealogical discordance in the group. Taken together these results suggest that there is a large amount of topological discordance among species in the personate and tubular clade, but there is generally support for monophyly at the species level.

### *Penstemon* subsect. Penstemon species exhibit variation in floral occlusion and pleat depth

Our quantitative morphological analyses confirm that the sampled subsect. *Penstemon* species vary in occlusion - encompassing a gradient of floral shapes from personate to open (Figure 3; Table 1). The three species described as personate have more occluded flowers than tubular species, which have more occluded flowers than open species (Figure 3). This result is significant when accounting for phylogenetic relatedness (*F* = 59.28932, *p* = 0.001; Appendix S7). Therefore, the qualitative floral descriptions by Pennell (1935) and Freeman (2019) capture quantifiable differences in floral shape. We also found a strong association between the degree of flower occlusion and pleat depth (slope = 2.238237, *p* < 0.0001; Appendix S8), further supporting the important role ventral floral pleats play in producing *Penstemon*’s personate flowers. Interestingly, such ventral floral pleats are absent from the corolla tubes of snapdragon (Appendix S3), suggesting convergent evolution of personate flowers in the two plant genera involves distinct structural mechanisms.

**Figure 3.**
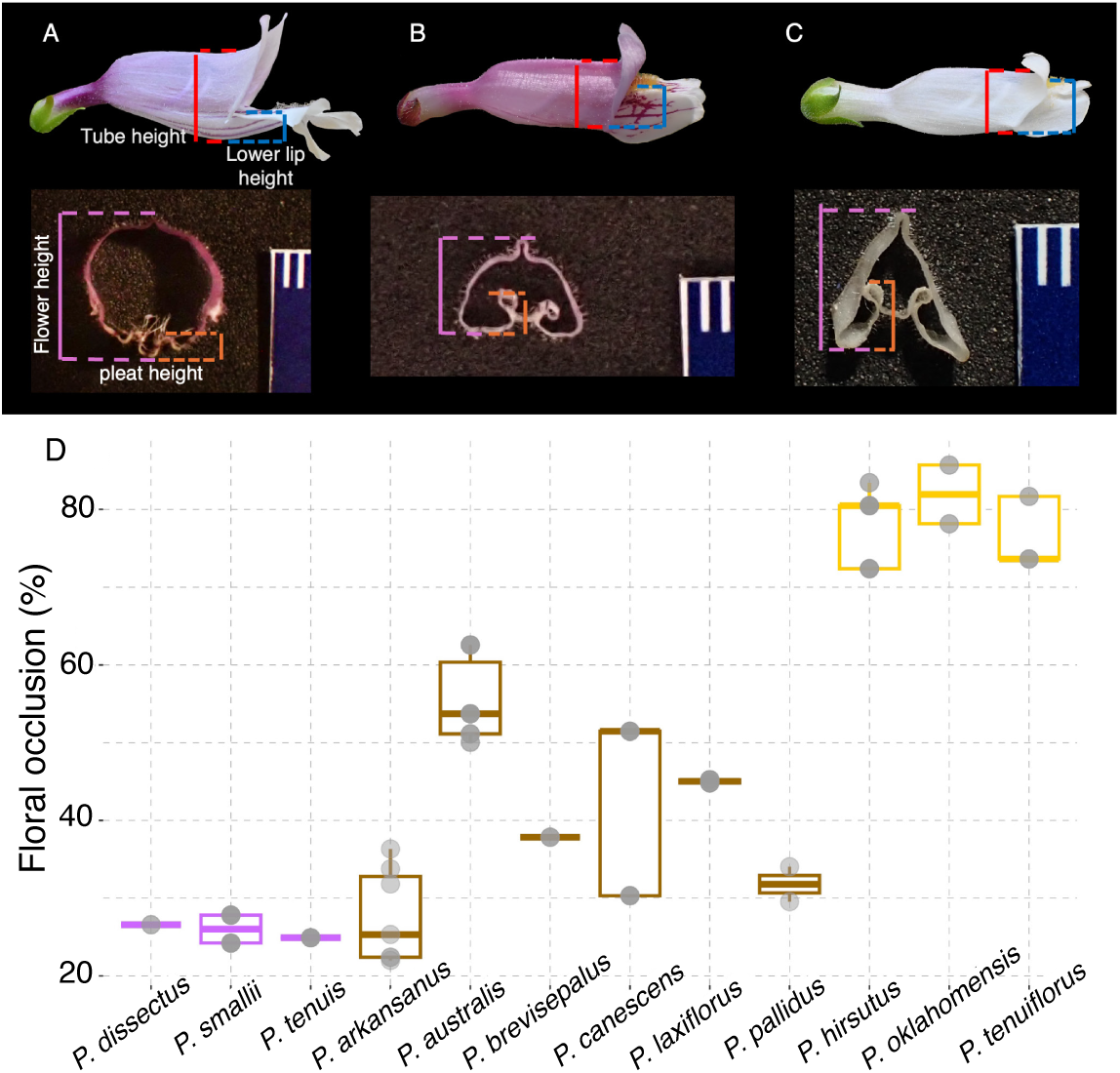
Measurements to quantify flower occlusion from lateral photographs and pleat depth from cross-section photographs. (A) *P. smallii*, an open-tubed species. (B) *P. australis,* a tubular species. (C) *P. hirsutus*, a personate species. (D) Variation in floral occlusion across several subsect. *Penstemon* species. Box outline color indicates the floral morphology of each species (purple: open, brown: tubular, and yellow: personate) and points represent population averages.

### *f*-branch indicates that many eastern *Penstemon* species have a history of allele sharing

Although our phylogenetic analysis suggests multiple transitions to personate flowers in subsect. Penstemon, used an *f*-branch analysis to examine whether the personate lineages exhibit a history of allele sharing consistent with introgression. This analysis revealed significant allele sharing between lineages, suggesting a history of introgression in subsect. Penstemon (Figure 4). For example, we find a striking signature of allele sharing between a branch of two eastern tubular species (*P. canescens* and *P. brevisepalus*) and almost all other tubular and personate species in the tree, particularly with the eastern personate species (*P. hirsutus* and *P. tenuiflorus*). *P. tenuiflorus* exhibits allele sharing with multiple clades. We also see a history of allele sharing between western tubular and eastern personate species. Shared geographic ranges and an overlap in ecological niche is likely associated with the hybridization signal among most *Penstemon* species. Interestingly, we do not find evidence of significant allele sharing between the eastern and western personate lineages, suggesting the two distinct personate lineages do not exhibit an elevated genome-wide signal of introgression.

**Figure 4.**
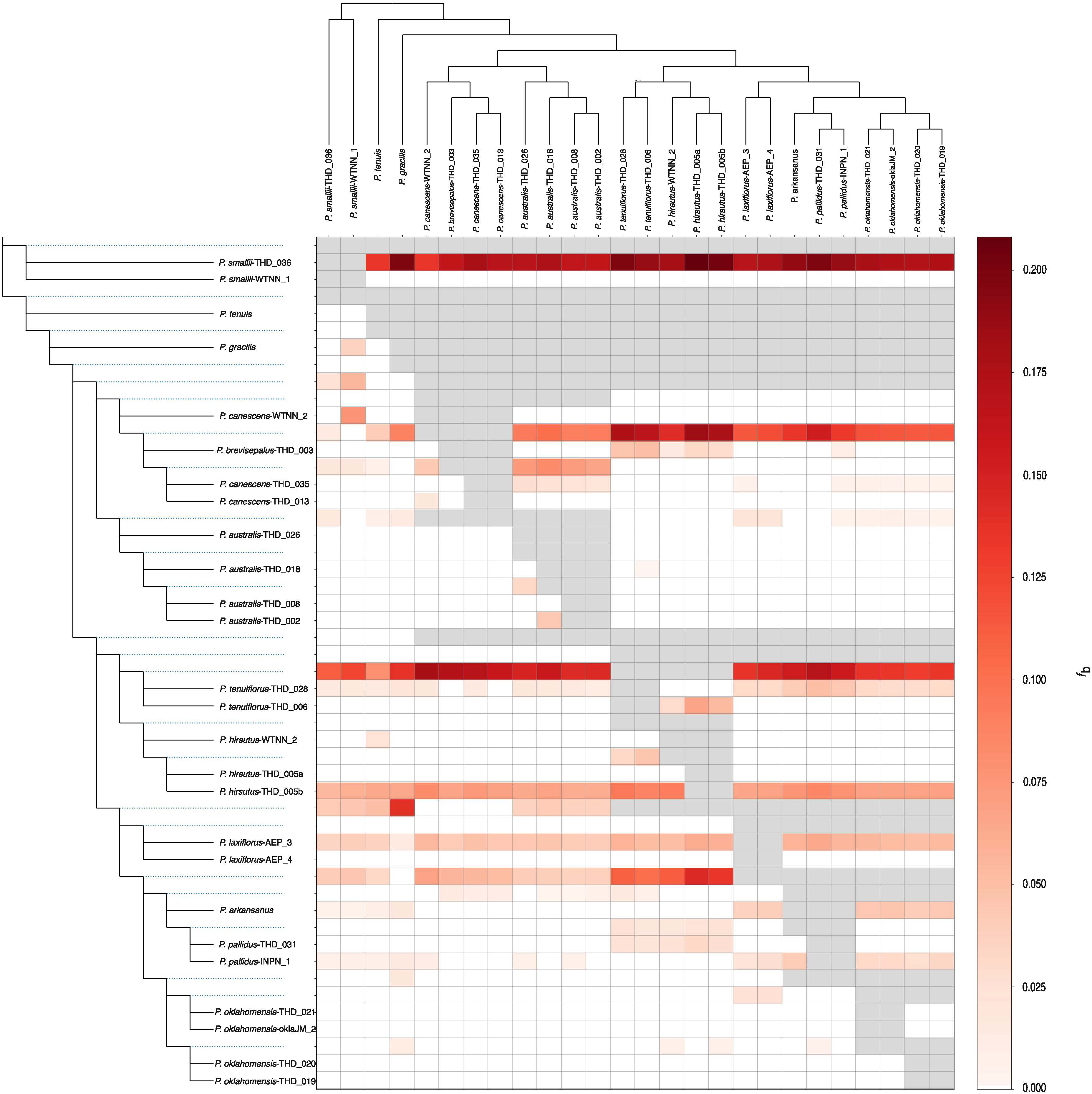
Patterns of introgression (*f*-branch) across the subsect. *Penstemon* species tree. The *f*-branch statistic (*f_b_*) is a genome-wide metric interpreted as excess allele sharing between branches *b* (y-axis) and tips (x-axis) in the species tree. The color gradient represents the *f_b_* score; gray boxes represent tests that are inconsistent with the species tree topology.

### Topology weighting reveals genealogical discordance and regions of the genome that support monophyly of personate taxa

Although we did not find genome-wide elevated patterns of allele-sharing between the eastern and western personate clades, there may be localized regions of the genome exhibiting such a pattern. Therefore, we used topology weighting to quantify the weightings of three possible topologies for the western personate, western tubular, and the eastern personate clades (Figure 5A). We found that most windows do not strongly support any of the three topologies suggesting an abundance of discordant site patterns and extensive ILS (Figure 5B). In our ternary plot, we found no left-right asymmetry in the distribution of topology weights, suggesting a similar chance that a given window tree resembles the two alternative topologies – again, consistent with ILS (Figure 5C). SNP windows that strongly support one of three topologies are dispersed across the genome (Figure 5D). Across the entire dataset of 78,057 100-SNP trees, we found that 11.5% of subtrees completely support the species tree (n=883), while 1.59% (n=122) completely support the miscellaneous discordant tree, and 1.37% (n=105) completely support the personate tree.

**Figure 5.**
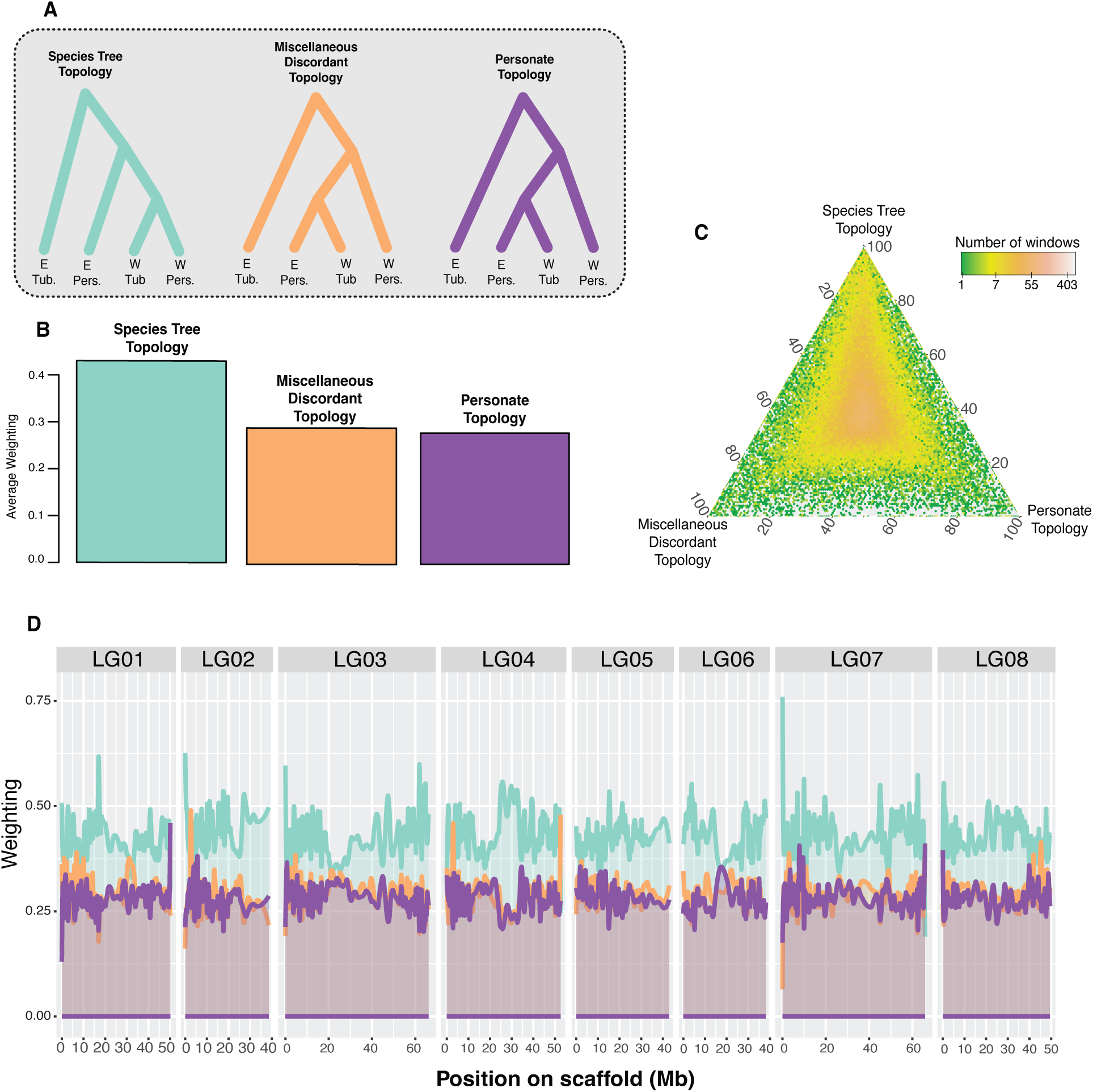
Topology weighting suggests ILS is an important contributor to phylogenomic discordance. (A) The three possible taxon topologies for the four taxa included in the analysis. (B) Genome-wide average weighting of each topology. (C) Ternary plot illustrating the distribution of topology weights for the 78,057 100-SNP windows. (D) Topology weightings for 100 SNP windows plotted across all eight linkage groups smoothed across 2 Mbp.

### Relative Node Depth analysis provides little evidence for a history of introgression between personate lineages

Our topology weighting analysis identified genomic windows that strongly support monophyly of personate species. Such discordant windows could reflect either ILS or introgression. Our relative node depth calculations are designed to examine these two possibilities. We expect ILS has greatly contributed to topological discordance in our data, and the distribution of RND between personate taxa (*RND_personate_*) in windows supporting the personate discordant topology should be similar (or shifted towards larger RND values) compared to the distribution of RND between the western tubular and western personate taxa (*RND_western_*) in windows supporting the species tree topology (Figure 6A). However, if introgression between the personate lineages has been an additional source of discordance, the distribution of *RND_personate_* values in the windows supporting the personate topology should show a second peak with a lower mean value, reflecting a relatively recent introgression event (Edelman et al., 2019).

**Figure 6.**
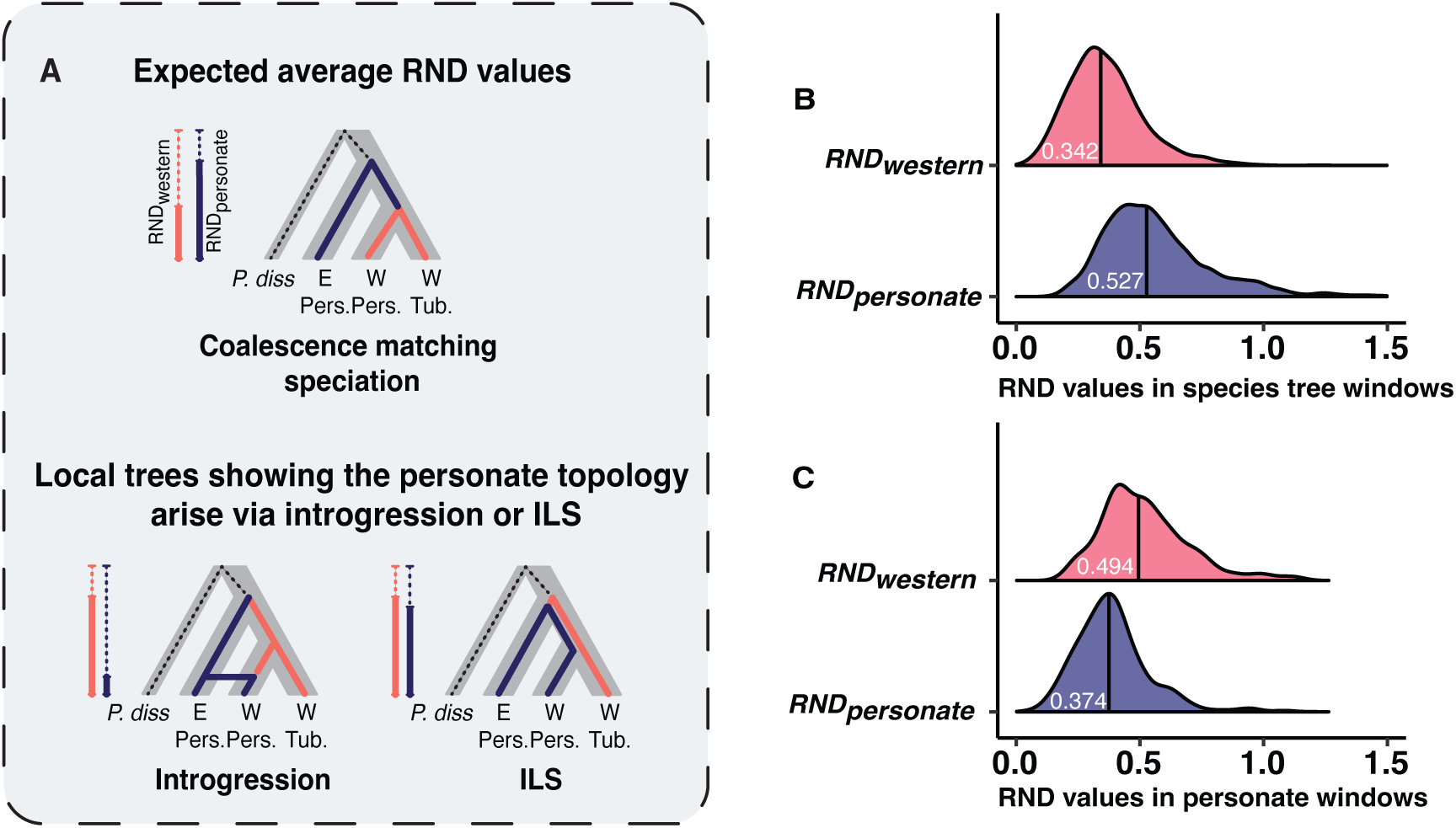
Relative Node Depth (RND) calculations suggest that ILS explains local windows that strongly support monophyly of personate lineages. (A) Hypothetical RND results under introgression and ILS. (B) Distribution of RND calculations in windows with a species topology weighting > 0.90. (C) Distribution of RND calculations in all windows with a personate topology weighting > 0.90. The black vertical line and white text represent the median RND value for each of the compared taxa (listed on y-axis).

We found that the distribution of *RND_personate_* values in windows that support the personate topology has a a single peak with median value that is greater than the distribution of *RND_western_* values in windows that support the species tree, suggesting most discordance supporting the personate topology is not due to post-speciation introgression (Figure 6B-C). The deep divergence in the majority of personate topology windows is instead consistent with ILS. However, there are some windows supporting the personate topology that have very small *RND_personate_* values, which might be consistent with introgression. This result suggests that ILS has been a common source of genealogical discordance causing local windows to show monophyly for the personate topology. We conducted an additional RND analysis for a more stringent set of windows (those fully supporting either the species tree or personate topologies) and found highly similar results (Appendix S9).

### Subsect. Penstemon species display a geographic pattern of relatedness

To examine whether population relatedness among tubular and personate species reflects geographic proximity, we summarized genomic variation using multi-dimensional scaling of pairwise genetic distance matrix as well as using a genomic PCA. We found a geographic pattern of relatedness among these taxa, perhaps reflecting a history of expansion eastwards – this pattern is present in both our MDS (Figure 7A) and our genomic PCA (Appendix S10). In fact, the MDS axis 1 and genomic PC1 both capture longitude, suggesting that populations and species are structured along a west-to-east gradient (Figure 7B-C; Appendix S10).

**Figure 7.**
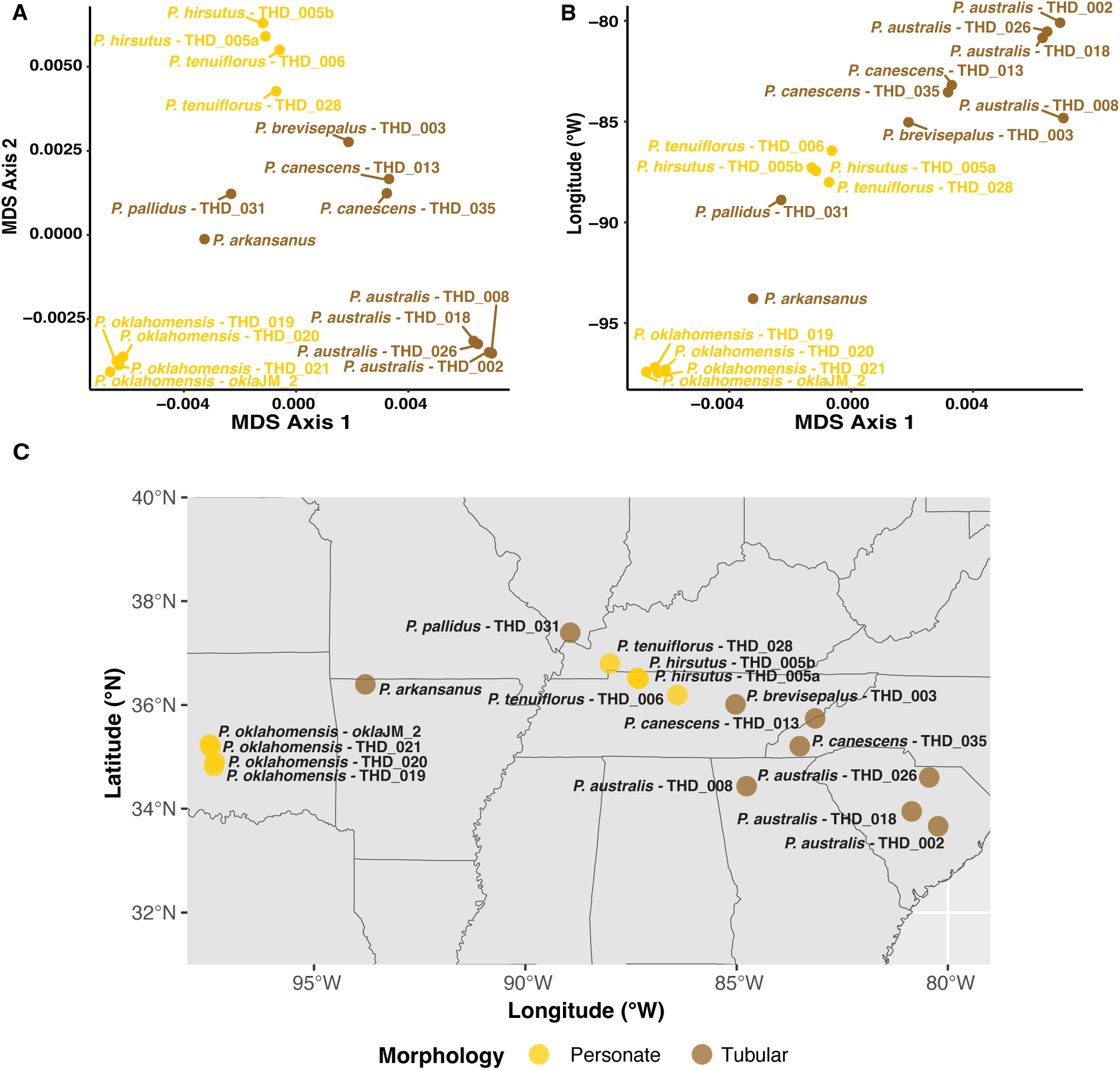
Population relatedness across subsect. Penstemon tubular and personate species mirrors geographic location. (A) Multi-Dimensional Scaling (MDS) plot for tubular and personate species. (B) Positive relationship between MDS Axis 1 and Longitude. Text and point colors represent the floral morphology of each species (brown: tubular and yellow: personate).

## DISCUSSION

### Personate flowers in *Penstemon* and *Antirrhinum* have convergently evolved through distinct structural mechanisms

*Penstemon*’s personate flowers are formed by pleats found on the ventral petal tube that project inward into the corolla tube (Figure 3A-C). These pleats shape the upward curvature of the ventral petal surface, causing occlusion of the floral tube. The ventral pleats are also present, but less pronounced, in open-tubed and tubular species. In fact, we found that floral occlusion in subsect. Penstemon is strongly predicted by pleat depth (Appendix S8). Since many bee-pollinated *Penstemon* species possess small ventral pleats, it is possible that the inward projections of floral tissue are a feature of *Penstemon* flower’s bilateral symmetry that support for the ventral petal lobe during insect visitation. However, we can speculate that the novel personate floral form involved co-option and deepening of these ventral pleats.

Snapdragon flowers lack these ventral petal pleats and instead achieve their fully occluded personate shape with a hinge structure which is formed where the upper and lower petal lobes meet (Appendix S3). Thus, personate flowers have convergently evolved across genera through different morphological structures.

### Pollinator-mediated selection and the evolution of personate flowers

In snapdragons, personate flowers are associated with bee pollination. Guzman and colleagues (2015) surveyed the morphologically diverse tribe Antirrhineae (28 genera), to examine whether the evolution of personate flowers is shaped by pollinator-mediated selection. The authors found that bees are the predominant visitor of species with personate flowers, however, other insect groups or even hummingbirds were occasionally reported as floral visitors suggesting that the personate flower is not always effective at excluding curious or persistent visitors. The strength of snapdragon’s floral hinge led to the hypothesis that only large bees were strong or heavy enough to pollinate the flowers (Müller, 1929; Sutton, 1988). According to this hypothesis, personate flowers might be an adaptation to prevent visitation by small bees that might be less effective pollinators. Vargas et al. (2010) tested this hypothesis by performing pollinator observations for three species snapdragons, finding that both large and small bees could visit the flowers, but flowers visited by large bees had the highest reproductive success. This result suggests that in this study, large bees were more effective pollinators than small bees.

Personate flowers in *Penstemon* have also been hypothesized to act as a “size-filter” on bees. Pennell (1935) speculated that large bees are the pollinators of personate *Penstemon* species, as observed in snapdragon. However, decades later Crosswhite and Crosswhite (1966) proposed, based on unpublished personal observations, that small bees, specifically those from the genera *Hoplitis* (Megachilidae) and *Ceratina* (Ceratinidae), are the primary pollinators of *P. hirsutus* and likely *P. tenuiflorus*. Clements et al. (1999) published the only study reporting pollinator observations in personate *Penstemon*, finding that two species of large bumblebee (*Bombus pennsylvanicus* and *B. bimaculatus*) were the primary pollinators of *P. hirsutus* and *P. tenuiflorus*. These conflicting hypotheses and scant empirical data reinforce the need for a comprehensive pollination study of the personate *Penstemon* species that includes both visitation and effective pollination data. Such data will inform on whether pollinator-mediated selection pressures influenced the evolution of personate flowers in *Penstemon*.

One intriguing difference between the evolution of personate flowers in *Penstemon* and snapdragons is in their associated flower colors. Snapdragons and other personate species in the Antirrhineae tribe are brightly colored, ranging in hues from pink to yellow (Whibley et al., 2006; Ellis and Field, 2016). All three personate species in subsect. Penstemon exhibit strikingly similar suite of correlated floral traits, including a lack floral pigmentation and nectar guides, but having a hairy yellow staminode. The existence of this “personate syndrome” suggests the action of similar ecological pressures causing a pattern of correlational selection that may be distinct from selective pressures acting on personate flowers of Antirrhineae.

### The source of genetic variation for parallel evolution

Our phylogenomic analyses revealed that personate species in subsect. Penstemon are not monophyletic. Taken at face value, this result suggests that there have been two evolutionary transitions to personate flowers. There are three possible sources of genetic variation for such repeated evolution: de novo mutation, recurrent adaptation from standing genetic variation, or adaptive introgression. Adaptive introgression in particular is a compelling mechanism for the repeated evolution of complex adaptations such as personate flowers and their correlated suite of traits. For example, if the loci responsible for the personate flower syndrome are genetically linked, introgression might transfer multiple linked loci during an introgression event. In this scenario, we expect to see an extended genomic signal of allele sharing between the personate taxa reflecting a genomic block that introgressed from one personate lineage into another at some point in evolutionary time.

Our genomic analyses were designed to test for a signal of introgression between the two personate lineages. Our genome-wide *f*-branch analysis found no evidence for a prevailing signal of introgression between personate taxa, nor did our topology weighting analysis find any extended blocks of allele sharing between the personate species. In contrast to our results, several other studies, for example in *Heliconius* butterflies (Martin and Van Belleghem, 2017; Rosser et al., 2024) and house mice (Linnenbrink et al., 2020) have identified large genomic regions exhibiting a signature of introgression using topology weighting. These genomic regions were found to harbor genes of adaptive significance such as the well-studied wing patterning gene - *optix* - in *Heliconius* butterflies (Pardo-Diaz, 2012).

If introgression between personate lineages has occurred in subsect. Penstemon, it did not leave an extended genomic signature. Nonetheless, it is possible that introgressed regions have a sufficiently small footprint that it is not detectable in the genome. Our *Twisst* analysis did detect regions of the genome where personate species are monophyletic, although they are relatively few and scattered throughout the genome. These local genomic regions showed relatively deep divergence times between personate lineages, pointing towards ILS rather than introgression as the source of discordant topologies showing monophyly of personate species. We conclude that introgression between the personate-flowered lineages has been comparatively rare compared to the overall history of introgression in subsect. Penstemon. We do not think it is likely that repeated shifts to personate flowers in this group has involved introgressed alleles, yet we cannot rule out the possibility completely.

Differentiating between the potential sources of genetic variation is often only possible once the causal genes have been identified. Within flowering plants, decades of research have focused on identifying the molecular pathways that produce flower pigments (Holton and Cornish, 1995; Dogbo et al., 1998), enabling the source of genetic variation for adaptation to be identified in diverse systems, including *Penstemon*. For example, functional genetic studies found that the evolution of red flowers in *Penstemon* appear to involve de novo loss-of-function mutations to the anthocyanin pathway gene *Flavonoid 3’, 5’-hydroxylase* (*F3’5’h*; Wessinger and Rausher 2014, 2015). The identification of this locus as a key component of floral syndrome divergence provides the opportunity for additional studies in *Penstemon* to examine the genealogy at this locus compared to the genome-wide phylogeny. Stone and Wessinger (2024) focused on a subgenus of *Penstemon* (Dasanthera) that includes two species that display a general hummingbird-pollination syndrome to identify the source of genetic variation (i.e. introgression, standing genetic variation, or de novo mutation) for the repeated evolution of bright magenta flowers. Similar to results reported here, this study found evidence that the two separate origins of magenta flowers involved distinct de novo mutations to *F3’5’h*.

To disentangle the source of genetic variation for separate transitions to personate flowers in subsect. Penstemon system we first need to identify the loci responsible for the personate syndrome traits. Prior work in the snapdragon model system using genetic manipulations have revealed the involvement of several symmetry genes (as reviewed in Hileman, 2014), boundary genes (Rebocho et al., 2017b) and differential growth patterns across epidermal flower cells (Coen and Rebocho, 2016; Rebocho et al., 2017a), suggesting there is a complex developmental genetic basis for this morphological trait. We currently lack information regarding the genetic basis of personate flowers in *Penstemon*, however, the occurrence of close relatives in sect. Penstemon with either personate or open flowers makes genetic mapping studies feasible. Identifying the genetic architecture of personate flowers in *Penstemon* will not only provide an opportunity to understand whether the same loci are responsible for personate flowers across the genus but will also allow us to examine the evolutionary history of personate flower alleles and ultimately determine whether introgression, standing variation or de novo mutation influenced the repeated evolution of this enigmatic floral type.

### Genealogical discordance in subsect. Penstemon reflects a rapid evolutionary radiation eastwards

We found evidence for substantial phylogenomic discordance in our sampled eastern *Penstemon* species. Our concordance factors indicate that loci exhibit conflicting evolutionary histories which is especially true in the clade with tubular and personate species (Figure 2). The phylogenomic discordance seen in our study is consistent with prior phylogenomic studies in genus *Penstemon* (Wessinger et al. 2019; Wolfe et al. 2021; Stone and Wessinger, 2024). The extensive genomic discordance in *Penstemon* can likely be attributed to due to its young age (about 2.5 million years old) and rapid species diversification across North America (Wolfe et al., 2021). We also identified a geographic pattern of genetic relatedness within subsect. Penstemon. This clade is inferred to be the youngest section of *Penstemon* - 1 million to 0.5 million years old (Wolfe et al., 2021) and many eastern species have partially overlapped ranges with similar ecological niches (Pennell, 1935; Freeman, 2019). Adaptive radiations often diversify quickly leaving porous barriers between closely related taxa in sympatry (Seehausen, 2004). The geographic pattern of relatedness within this group likely reflects its recent geographic expansion eastwards.

## CONCLUSIONS

Personate flowers represent a complex morphological innovation with a unique structural basis in *Penstemon* that apparently has evolved on a short evolutionary timescale. However, we currently have information on how and why an ancestrally open-tubed lineage evolves personate flowers. Future research should investigate the genetic architecture underlying this intriguing floral trait to determine how many loci are involved, the developmental genetic pathways responsible, and whether the repeated evolution of personate flowers involves de novo mutations or pre-existing variants. Additionally, there is a pressing need to identify the visitors and effective pollinators of the subsect. Penstemon species. Such studies will shed light on potential pollinator-mediated selection pressures that may have favored this unique floral shape in *Penstemon*.

## AUTHOR CONTRIBUTIONS

T.H.D. and C.A.W. conceived the study, T.H.D. collected and analyzed the data, T.H.D and C.A.W. wrote the manuscript.

## ACKNOWLEDGEMENTS

We thank B. Stone for advice on phylogenomic analyses and comments on an initial version of this manuscript. We thank Phase Genomics for assembling and annotation of the *P. smallii* genome. We thank C. Bellinger, A. Hamilton, and J. Stevens for assistance collecting plant specimens used in this study. We are grateful for C. Freeman who provided samples of *P. arkansanus* and *P. gracilis* from the R. L. McGregor Herbarium at the University of Kansas. We thank J. Messick for providing species information and a leaf sample of *P. oklahomensis*. We appreciate the iNaturalist users Boverser, Brian Finzel, Jared Gorrell, Luke Benjamin, and Theo Witsell for allowing us to use their photos of *P. arkansanus* and *P. pallidus* in this study. We are grateful to the USDA Forest Service collection permits in the Nantahala and Pisgah National Forests (NC), the Georgia Department of Natural Resources for collection permits for *P. dissectus*, and the Southeastern Climbers Coalition for allowing us to collect *P. hirsutus* at Kings Bluff (Clarksville, TN). This work was funded by NIH NIGMS R35GM142636 (to C.A.W.) and NSF DEB-2052904 (to C.A.W.).

## DATA AVAILABILITY STATEMENT

All raw genomic data and the annotated *P. smallii* assembly are accessible under NCBI BioProject PRJNA1188554. The data that support the findings of this study will be deposited in publicly accessible databases upon acceptance of this manuscript.

## SUPPORTING INFORMATION

**Appendix S1.** Voucher information for *Penstemon* samples used in this study.

**Appendix S2.** Information for diploid species of *Penstemon* subsect. Penstemon used in this study from Pennell (1935) and The Flora of North America (Freedman, 2019).

**Appendix S3.** Photos demonstrating snapdragon (*Antirrhinum*) flower morphology.

**Appendix S4.** Density of physical size (kbp) of non-overlapping windows of 100 SNPs used for *Twisst* and RND analyses.

**Appendix S5.** Species tree constructed in Astral-III from 20kb windows.

**Appendix S6.** Maximum Likelihood concatenated species tree constructed in IQ-TREE.

**Appendix S7.** Floral occlusion phylogenetic ANOVA statistical information.

**Appendix S8.** Comparison between the results of two methods developed to quantify eastern *Penstemon* flower morphology (floral occlusion and pleat depth).

**Appendix S9.** Relative Node Depth (RND) calculations to disentangle the genomic signatures of introgression and ILS within the miscellaneous discordant *Twisst* topology.

**Appendix S10.** Relationship between geography and genomic PCA.

