## Supporting Material for "The unique morphological basis and parallel evolutionary history of personate flowers in *Penstemon*"

**Appendix S1.** Voucher information for *Penstemon* samples used in this study. All vouchers deposited in R.L. McGregor Herbarium (KANU) and/or A.C. Moore Herbarium (USCH) unless otherwise noted.

| **Taxon** | **Voucher information** | **Herbarium information** |
|---|---|---|
| *P. arkansanus* | Craig C. Freeman #25396 | KANU |
| *P. australis* | THD_002 | KANU |
| *P. australis* | THD_008 | KANU |
| *P. australis* | THD_018 | KANU, USCH |
| *P. australis* | THD_026 | Photo Voucher |
| *P. brevisepalus* | THD_003 | KANU |
| *P. canescens* | canescens2_WTNN | Photo Voucher |
| *P. canescens* | THD_013 | KANU |
| *P. canescens* | THD_035 | USCH |
| *P. dissectus* | THD_010 | KANU |
| *P. gracilis* | Craig C. Freeman #24662 | KANU |
| *P. hirsutus* | hirsutus2_WTNN | Photo Voucher |
| *P. hirsutus* | THD_005.21a | USCH |
| *P. hirsutus* | THD_005.22b | USCH |
| *P. laxiflorus* | laxiflorus3_AEP | Photo Voucher |
| *P. laxiflorus* | laxiflorus4_AEP | Photo Voucher |
| *P. oklahomensis* | oklaJM_2 | Photo Voucher |
| *P. oklahomensis* | THD_019 | KANU |
| *P. oklahomensis* | THD_020 | KANU, USCH |
| *P. oklahomensis* | THD_021 | KANU, USCH |
| *P. pallidus* | pallidus1_INPN | Photo Voucher |
| *P. pallidus* | THD_031 | Photo Voucher |
| *P. smallii* | smallii1_WTNN | Photo Voucher |
| *P. smallii* | THD_036 | KANU |
| *P. tenuis* | tenuis3_AEP | Photo Voucher |
| *P. tenuiflorus* | THD_006 | KANU |
| *P. tenuiflorus* | THD_028 | KANU |

**Appendix S2.** Information for diploid species of *Penstemon* subsect. Penstemon used in this study from The Scrophulariaceae of Eastern Temperate North America (Pennell, 1935) and The Flora of North America (Freedman, 2019).

| Species name | Flower morphology | Flower color | Nectar guides | Geographic description |
|---|---|---|---|---|
| *P. dissectus* | Open | Lavender to violet-purple | Violet-purple | Coastal plain of Georgia |
| *P. smallii* | Open | Lavender to violet-purple | Violet-purple | Southern Appalachian Mountains |
| *P. tenuis* | Open | Lavender to violet-purple | Reddish to violet-purple | Widespread in Western Gulf coast plain and lower Mississippi River basin |
| *P. gracilis* | Tubular | Pale lavender to violet-purple | Violet-purple | Widespread in grass prairies and foothills of Northern United States |
| *P. canescens* | Tubular | Pale lavender to violet-purple | Lavender or violet-purple | Widespread East of the Mississippi River |
| *P. brevisepalus* | Tubular | Pale lavender to purple | Light purple | Range lies between *canescens* and *pallidus* |
| *P. australis* | Tubular | Dark pinkish red to purple | Dark reddish to purple | Southern Atlantic coastal plain and Eastern Gulf coast plain |
| *P. laxiflorus* | Tubular | White to light lavender and sometimes tinted pink | Reddish-purple | Central and Western Gulf coast plain and Southern interior lowland |
| *P. arkansanus* | Tubular | Corolla white sometimes tinted pinkish | Reddish to violet-purple | Interior highlands |
| *P. pallidus* | Tubular | Corolla white sometimes tinted lavender-purple | Reddish to violet-purple | Central lowlands and Ozark plateau |
| *P. hirsutus* | Personate | Corolla white or slightly violet tinged externally | None | Widespread East of the Mississippi River |
| *P. tenuiflorus* | Personate | Corolla white or slightly lavender or purplish | None | Interior low plateaus region of Western Kentucky, central Tennessee, North-West Alabama, Black-belt section of central Alabama, and North-East Mississippi |
| *P. oklahomensis* | Personate | Corolla white to ochroleucus (yellowish-white) | None | Oklahoma and Northern Texas |

**Appendix S3.** Photos demonstrating snapdragon (*Antirrhinum*) flower morphology. (A) lateral flower (B) lateral flower with petal lobes removed (C) smooth underside (ventral) of the flower (D) lower petal lobe separated from the rest of the flower (E) snapdragon cross-section. Blue arrow represents location of the floral hinge, orange arrow highlights the lower petal lobe ridges. All photos by T. Depatie.

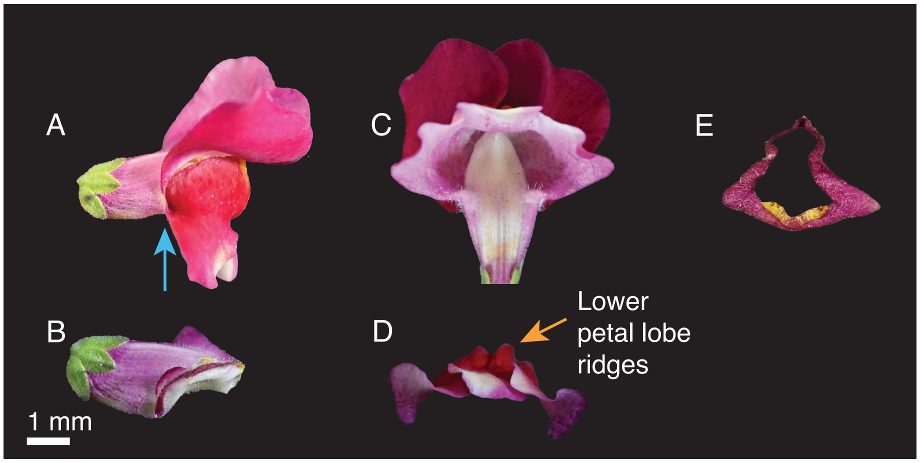

**Appendix S4.** Density of physical size (kbp) of non-overlapping windows of 100 SNPs used for *Twisst* and RND. The median window size, indicated by the vertical line, is 2.1kb.

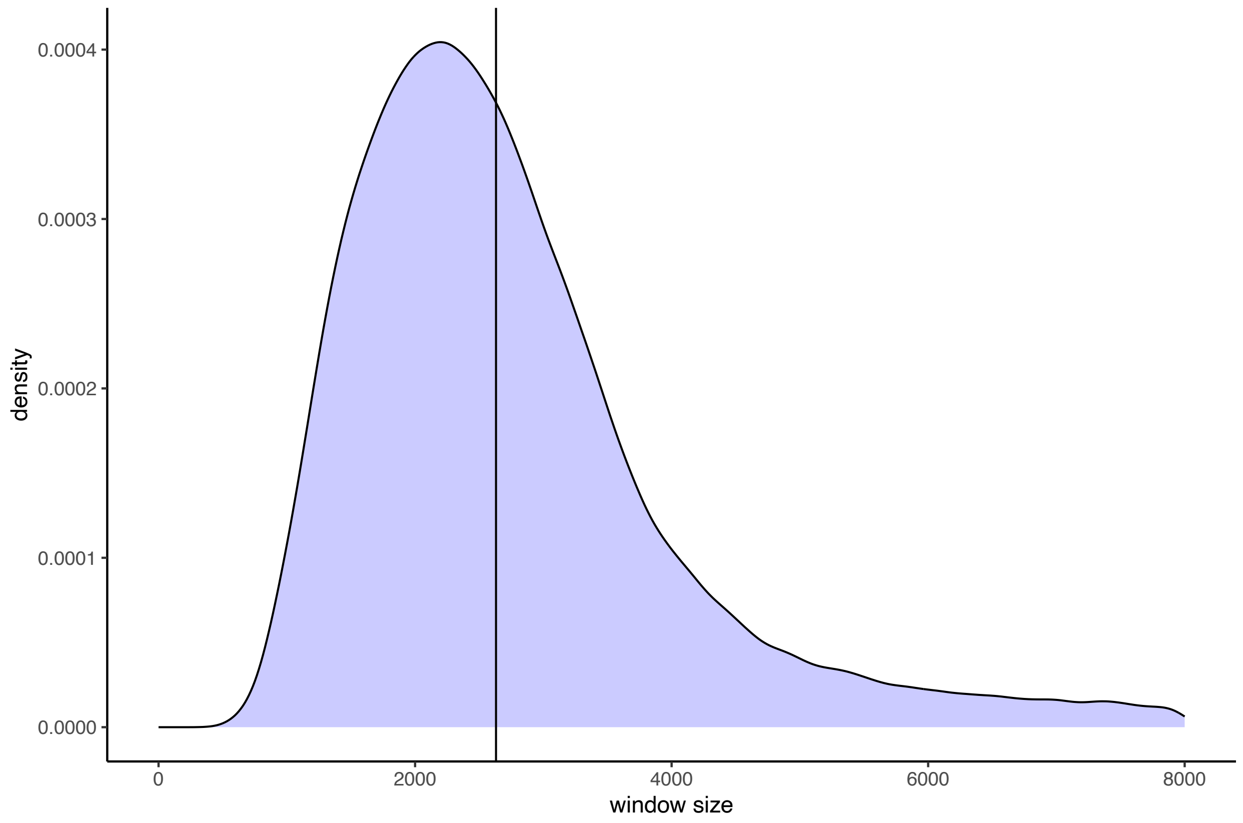

**Appendix S5.** Species tree constructed in Astral-III from 20kb windows.

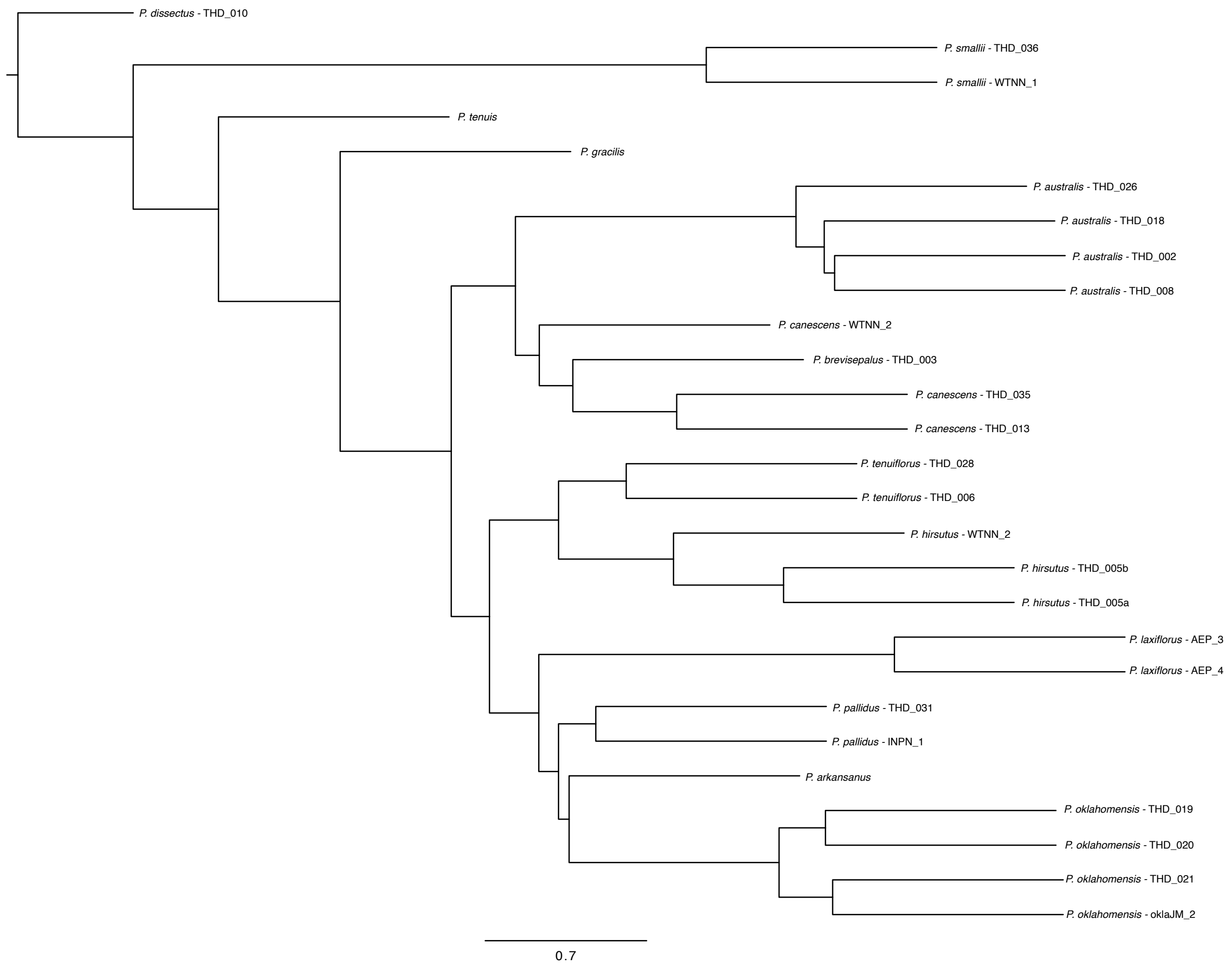

**Appendix S6.** Maximum Likelihood concatenated species tree constructed in IQ-TREE.

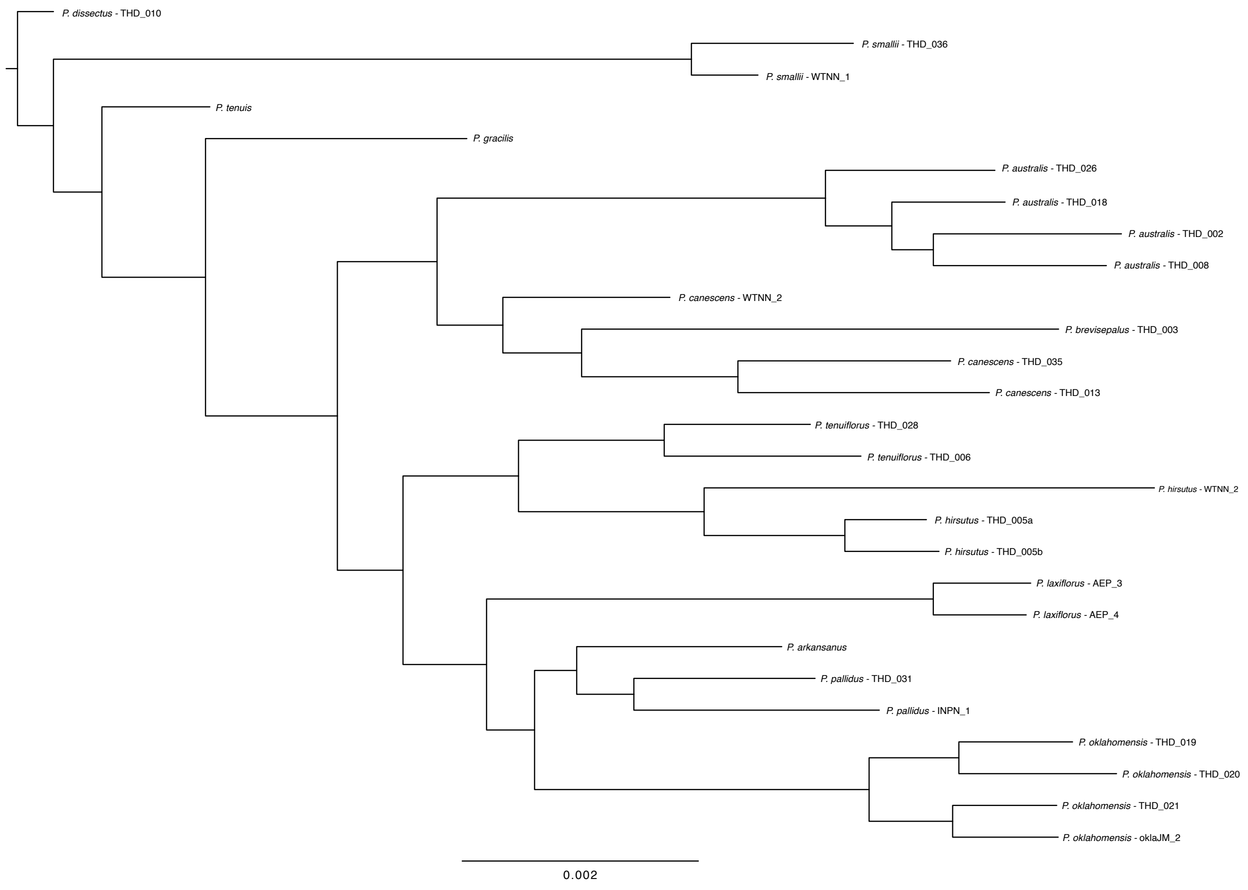

**Appendix S7.** Floral occlusion phylogenetic ANOVA statistical information. Pairwise posthoc test was completed with the "holm" method. Pairwise t-values are reported here since they are the difference between the means of each flower morphology group.

**Pairwise t-values:**

| | **Open** | **Personate** | **Tubular** |
|---|---|---|---|
| **Open** | 0.000000 | -10.000062 | -3.683654 |
| **Personate** | 10.000062 | 0.000000 | 8.515269 |
| **Tubular** | 3.683654 | -8.515269 | 0.000000 |

**Pairwise corrected P-values:**

| | **Open** | **Personate** | **Tubular** |
|---|---|---|---|
| **Open** | 1.000 | 0.003 | 0.032 |
| **Personate** | 0.003 | 1.000 | 0.003 |
| **Tubular** | 0.032 | 0.003 | 1.000 |

**Appendix S8.** Comparison between the results of two methods developed to quantify eastern *Penstemon* flower morphology (floral occlusion and pleat depth).

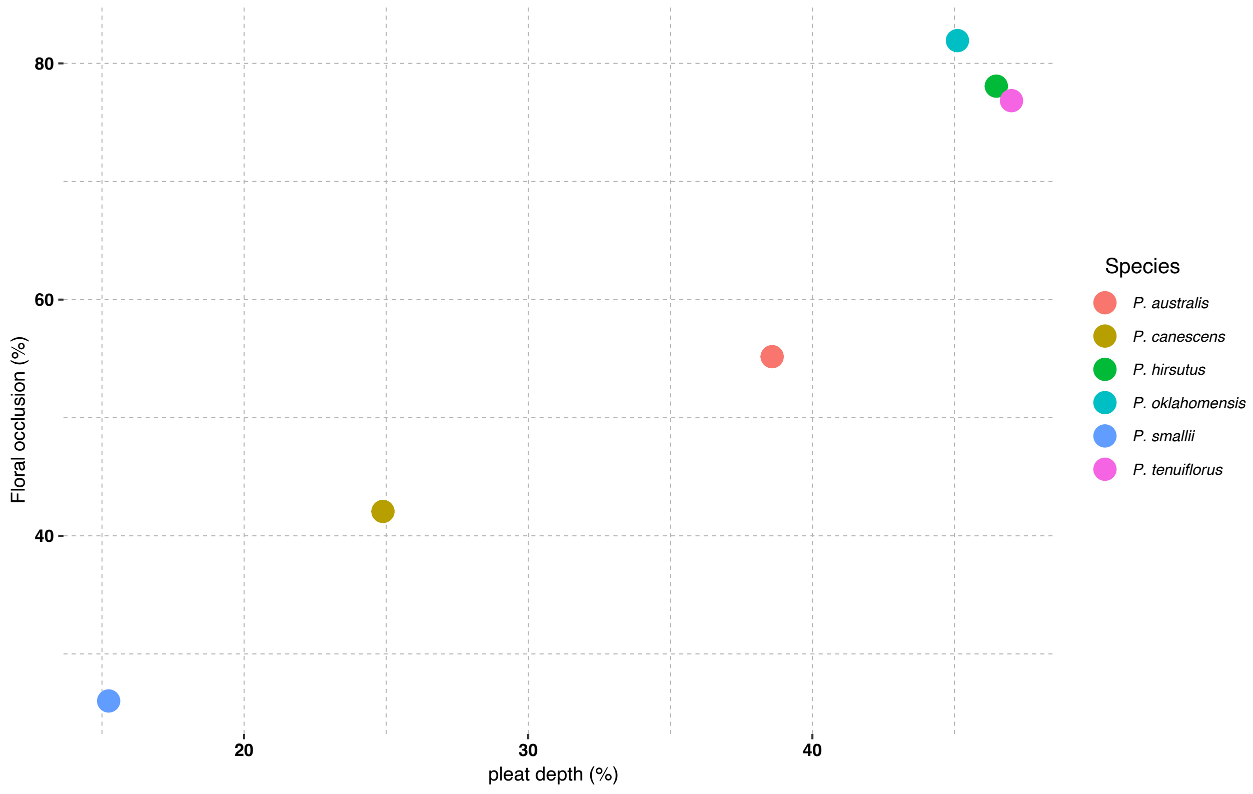

**Appendix S9.** Relative Node Depth (RND) calculations suggest that ILS explains local windows that strongly support monophyly of personate lineages. (A) Hypothetical RND results under introgression and ILS. (B) Distribution of RND calculations in windows fully supporting the species topology (weighting of 1.0). (C) Distribution of RND calculations in windows fully supporting the personate topology (weighting of 1.0). The black vertical line and white text represent the median RND value for each of the compared taxa (listed on y-axis).

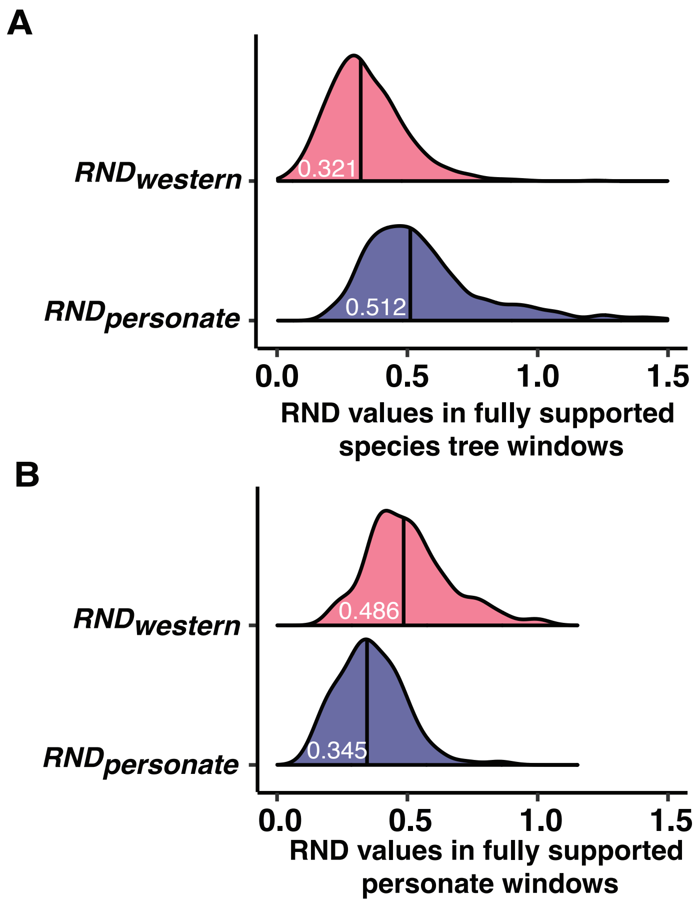

**Appendix S10.** Relationship between geography and phylogenetic relatedness. (A) Genomic PCA for tubular and personate species and (B) strong correlation between PC1 and Longitude. Text color indicates the floral type of each species (brown: tubular and yellow: personate). Open species were excluded from these analyses.

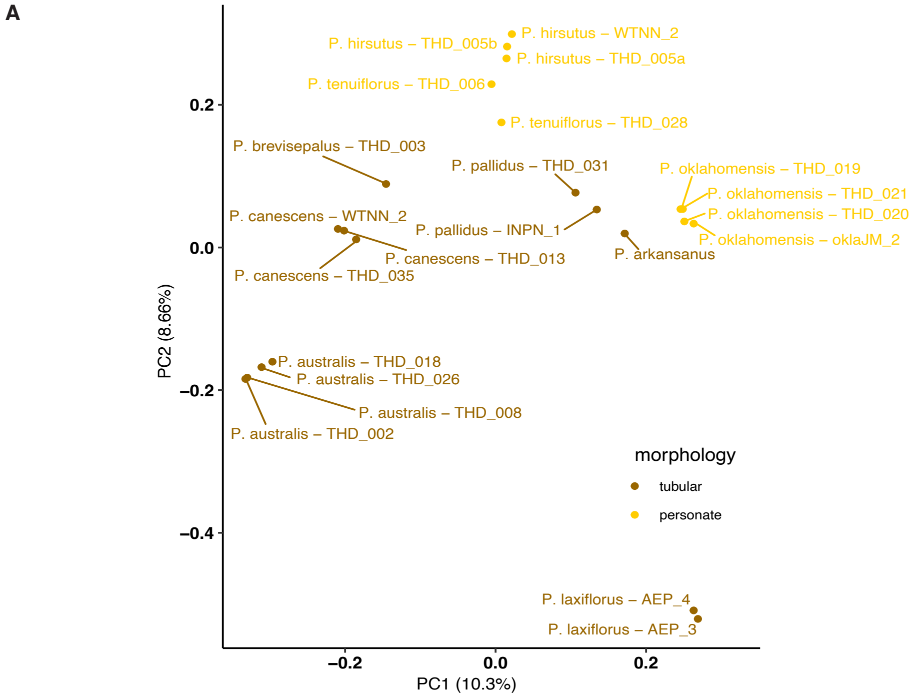


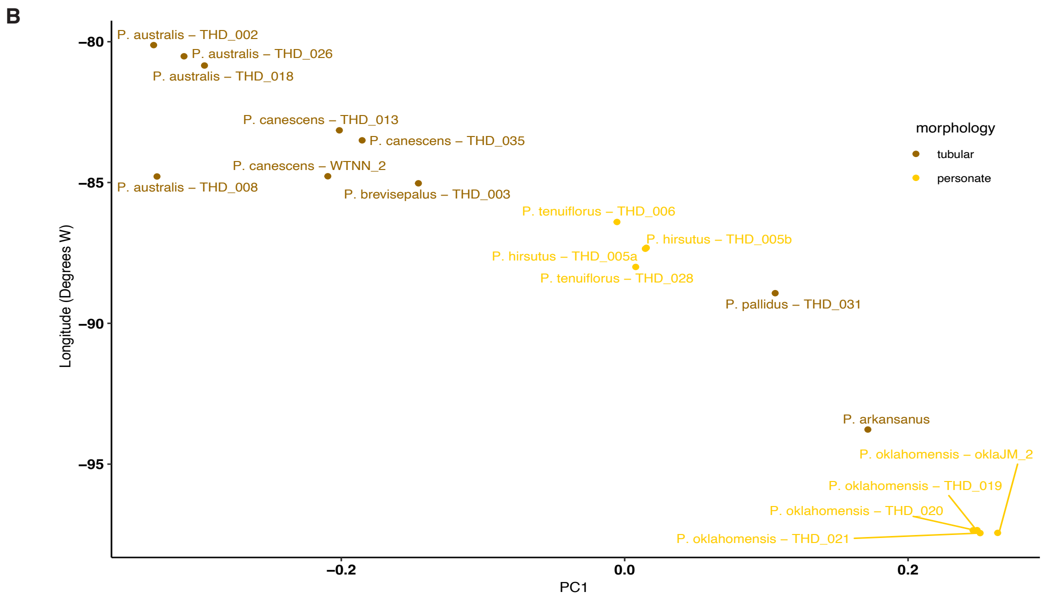
